# Functional Analysis of Enzyme Families Using Residue-Residue Coevolution Similarity Networks

**DOI:** 10.1101/646539

**Authors:** Christian Atallah, David James Skelton, Simon J. Charnock, Anil Wipat

## Abstract

**Motivation:** Residue-residue coevolution has been used to elucidate structural information of enzymes. Networks of coevolution patterns have also been analyzed to discover residues important for the function of individual enzymes. In this work, we take advantage of the functional importance of coevolving residues to perform network-based clustering of subsets of enzyme families based on similarities of their coevolution patterns, or “Coevolution Similarity Networks”. The power of these networks in the functional analysis of sets of enzymes is explored in detail, using Sequence Similarity Networks as a benchmark.

**Results:** A novel method to produce protein-protein networks showing the similarity between proteins based on the matches in the patterns of their intra-residue residue coevolution is described. The properties of these co-evolution similarity networks (CSNs) was then explored, especially in comparison to widely used sequence similarity networks (SSNs). We focused on the predictive power of CSNs and SSNs for the annotation of enzyme substrate specificity in the form of Enzyme Commission (EC) numbers using a label propagation approach. A method for systematically defining the threshold necessary to produce the optimally predictive CSNs and SSNs is described. Our data shows that, for the two protein families we analyse, CSNs show higher predictive power for the reannotation of substrate specificity for previously annotated enzymes retrieved from Swissprot. A topological analysis of both CSNs and SSNs revealed core similarities in the structure, topology and annotation distribution but also reveals a subset of nodes and edges that are unique to each network type, highlighting their complementarity. Overall, we propose CSNs as a new method for analysing the function enzyme families that complements, and offers advantages to, other network based methods for protein family analysis.

**Availability:** Source code available on request.

## 1 Introduction

Performing functional analysis of protein families and classifying their members is a major challenge in bioinformatics. For example, assigning functional annotations to enzyme families can lead to a better understanding of enzyme evolution and enzyme diversity (Baier *et al.* (2016)), with applications in areas such as enzyme engineering (Brown and Babbitt (2014)). Enzymes are also being increasingly used as biocatalysts in different fields such as the pharmaceutical industry, removing much of the need for expensive and environmentally-harmful chemical reagents used in synthesis reactions (Ran *et al.* (2008)).

However, functional analysis and annotation of enzymes is difficult, especially the assignment of substrate specificity. Indeed, while the number of available protein and enzyme sequences has increased exponentially thanks to next-generation sequencing, the proportion of sequences which have known experimental annotation has lagged behind. For example, UniprotKB is a central hub of proteomic data that is separated into two subsets: Swiss-Prot, whose entries are manually curated, and TrEMBL, whose entries are not (Apweiler *et al.* (2004)). As of the 13/02/2019 release, TrEMBL stores over 140 million sequences, while Swiss-Prot only contains around 550 thousand (Uniprot Consortium (2019a,b)). Furthermore, only around 28% of the Swiss-Prot proteins have experimental evidence at the transcript or protein level, with around 69% inferred from homology.

Swiss-Prot’s latest release also contains 238,254 entries with “catalytic activity’ annotation i.e. enzymes with annotated functions. Only 10,921 (4.5%) of these entries have their catalytic activity sections backed up by the evidence tag “ECO_0000269”, which is used by UniprotKB to indicate annotation that is “manually curated information for which there is published experimental evidence”. Indeed, of Swiss-Prot enzymes, 179,784 (74.5%) have their catalytic activity assigned by what Uniprot terms “sequence models”, which are generated by an automatic annotation system that uses predictive tools like HAMAP (Lima *et al.* (2009); Gattiker *et al.* (2003)). This system annotates entries by matching them to sets of template alignments and rules about important sequence features. While these templates are manually curated and kept up to date, sequence similarity is ultimately the core metric.

High sequence identity (>40%) has long been proven to be a good estimator for conserved protein structure and function, but it has significant drawbacks. The “twilight zone” (20-35 identity) is a well known threshold where sequence conservation stops being a useful metric for declaring two proteins as homologous (Rost (1999)). However, two enzymes lying in this “twilight zone” of uncertainty does not preclude them from being functionally similar (and vice versa) (Ponting (2017)). For example, lactonases exhibiting phosphotriesterase activity have been found in three different superfamilies: amidohydrolases, six-bladed propellers, and metallo-b-lactamases (Babtie *et al.* (2010)). Indeed, aligning sequences of example enzymes from each of these superfamilies (Q97VT7, P27169, A9CKY2, respectively) onto a multiple sequence alignment (MSA) results in percent identity values that range from 15.06% to 23.48% - figures which are far from the 40% threshold usually associated with homology (Rost (1999)). Clearly, new approaches are required to confirm functional similarity of proteins with sequence similarity that falls in and below the twightlight zone.

New approaches such as Sequence Similarity Networks (SSN) have been shown to be useful for giving a good overview of the functional diversity of enzyme families and superfamilies (Atkinson *et al.* (2009); Gerlt *et al.* (2015)), techniques that can overcome the aforementioned drawbacks associated with pure sequence similarity are needed.

Residue-residue coevolution is another useful method for searching for functional conservation in protein sequence and structure. Two amino acid residues are said to coevolve if an influential substitution in one residue is counteracted by a substitution in the other residue (Morcos *et al.* (2011)). Under an evolutionary lens, residue-residue coevolution in a protein can be seen as happening due to the evolutionary pressure on the protein for retaining functional stability. While residue-residue coevolution information has mostly been used to infer spatial constraints for residues for the purposes of tertiary structure prediction (Moult (2005); Moult *et al.* (1999); Schaarschmidt *et al.* (2018)) and for predicting protein-protein interactions (Pazos *et al.* (1997); Ovchinnikov *et al.* (2014)), patterns of coevolution have also been used in a functional context. Coevolution patterns are often represented as homogeneous residue-residue coevolution networks, where networks are proteins, nodes are residues, while edges between nodes are created if the two residues are likely to coevolve. Such networks have been successfully used in revealing functionally important residues (Salinas and Ranganathan (2018); Gloor *et al.* (2005); Yeang and Haussler (2007); Lee *et al.* (2008); Kuipers *et al.* (2009); Dwyer *et al.* (2013); Lee *et al.* (2012)). With the emergence of both better and faster methods for computing the coevolution metric on a larger scale (Hopf and Marks (2017); Ekeberg *et al.* (2013); Seemayer *et al.* (2014); Wang *et al.* (2017); Schaarschmidt *et al.* (2018)), it is now possible to produce better coevolution data for a set of enzymes and build residue-residue coevolution networks for each of them on a larger scale than previously possible.

While coevolution data has been used to analyse function in individual enzymes, to the authors knowledge, there has been little or no use for single protein co-evolution data as a similarity metric between proteins in a protein family-wide classification analysis. As coevolution patterns contain information important to function, the hypothesis explored in this work is that functionally similar enzymes, specifically in terms of their substrate specificity will share similar coevolution patterns. To tackle the aforementioned challenges of sequence similarity-based methods in functional analysis, we explored how an all-vs-all similarity comparison of the coevolution patterns of a set of enzymes could be used to perform a functional analysis of a dataset of coevolution patterns.

We also introduce a method of representing data about functionally related proteins that we call Coevolution Similarity Networks (CSN). CSNs are built using the same network logic as SSNs: nodes are proteins, and edges are made between nodes if their coevolution patterns are similar enough based on some set threshold.

In this work we show that CSNs can describe the distribution of functional diversity of a set of enzymes across a network, in terms of substrate specificity, in a way that groups functionally of similar nodes, in a similar, but complementary, fashion to SSNs. We compare the CSNs and SSNs produced for two different annotated enzyme datasets from Swiss-Prot (transaminases and short-chain dehyrodgenases) in terms of both the resulting networks that describe the family structure and in their predictive power in a label propagation experiment for predicting enzyme substrate specificity. We show that the CSN is able to predict known annotation of subsrate specifity more specifically than the SSN for these two examples. Finally also, consider the ability of CSNs to predict promiscious enzymes and show that CSNs could have role complementary to SSNs in the prediction of enzymes with multiple substrates.

## 2 Methods

### 2.1 The Datasets

To investigate the use of CSNs two enzymes families were used. The first comprises 241 transaminases, while the second is a set of 142 enzymes from the short-chain dehydrogenases/reductases (SDR) family. Both datasets were built from Swissprot (Apweiler *et al.* (2004)). These datasets were chosen since they are well annotated and relevant to the focus of this study and were built by searching on Swissprot for prokaryotic entries that contain the PFAM identifiers for the respective families, PF00202 for the transaminases and PF00106 for the SDRs (Bateman *et al.* (2004)). We refer to these datasets as Trans241 and SDR142 respectively for the remainder of this work.

The advantage of selecting datasets from Swissprot is that we can be more confident of their annotation. One important annotation type most of these entries contain is the Enzyme Commission (EC) number (Bairoch (2000)). The EC number is a hierarchical classification system that numerically labels an enzyme with four progressively more specific levels of functional detail, down to the reported substrate specificity.

An important drawback of EC numbers is that the specificity of an enzyme might be known, but not have a complete EC number as one for that substrate has not been curated yet. Also, some complete EC numbers will still be hierarchical in nature and correspond to entire classes of enzymes. For these reasons, we temporarily manually annotated the EC label of some entries (for example, where the substrate is known and cited in the Swissprot record), the list of which can be seen in Table S1. Tables S2 and S3 also contains the EC number distributions for both datasets.

### 2.2 SSN Construction

For both the Trans241 and SDR142 datasets, we produce SSNs throughout this work. We perform a global pairwise alignment using Needleman-Wunsch Needleman and Wunsch (1970) in an all-vs-all fashion, producing a sequence identity matrix for an entire dataset. This matrix is then used along with a threshold to build a SSN. We use a rigorous form of SSNs that require a global alignment algorithm with similarity over the whole length of the protein, whilst other methods typically used BLAST based approaches Atkinson *et al.* (2009).

### 2.3 Coevolution Preprocessing

We produced the coevolution data for every single protein using CCMpred (Seemayer *et al.* (2014)). Specifically, we followed the exact recommended steps for producing the data (https://github.com/soedinglab/CCMpred/wiki/FAQ). The result was a coevolution matrix for every individual enzyme in both datasets.

### 2.4 Residue-Residue Coevolution Network Construction

For each of the enzymes, the coevolution matrix was represented using NumPy (Walt *et al.* (2011)), and then the top N coevolving pairs were picked. As recommended by CCMPred, the top N pairs for each matrix where selected. In this work specifically, we picked N to be an arbitrarily high number to contain a high amount of coevolving pairs based on the average sequence length of the dataset (600 for the SDR142 dataset and 700 for the Trans241 dataset). From the coevolving pairs picked, we created residue-residue coevolution networks for each protein where nodes are residues and edges are made between the pairs that are said to coevolve.

### 2.5 Residue-Residue Coevolution Network Mapping

To compare the residue-residue networks, we needed a method of comparing a coevolving pair in one enzyme to a coevolving pair in another enzyme, and declaring them to be “equivalent”. To do this, we align the sequences into a MSA to establish the relative position co-evolving residues. We then assumed that if a coevolving pair of one enzyme aligns to a coevolving pair from another enzyme, then they are Equivalent Coevolving Pairs (ECP). We therefore used Clustal-Omega (Sievers *et al.* (2011)) to align all the sequences of a set into an MSA i.e. one MSA for the Trans241 dataset, and one MSA for the SDR142 dataset.

As a result, for each dataset, we create an intermediate single large alignment network where the nodes represent, collectively, the coevolving residues of all the residue-residue coevolving networks, and edges are made between nodes if they align on the MSA i.e. they are ECPs. For example, if residues 40-60 coevolve in A, and residues 42-62 coevolve in B, and the two pairs align on the MSA, then the two pairs are ECPs, and we make an edge between A-40 and B-42, and between A-60 and B-62.

It is important to note that for a given functional protein family some of the coevolving pairs are expected to be present in the majority of a families members (de Juan *et al.* (2013)). Coevolving pairs that are present in a majority of enzymes in a dataset are therefore uninformative when the purpose of the analysis is to discriminate function at the specificity level, where the enzyme class is common to all members under analysis. In these cases, we filtered out coevolving pairs from the alignment network that occur in an high proportion of the dataset members. Typically, coevolving pairs that occur in over 60% of a dataset were removed, as none of the enzyme substrate specificity classes in either datasets make up that large of a proportion. However, this parameter is variable and optional.

### 2.6 Clique-based Residue-Residue Coevolution Network Comparison

For each residue-residue coevolution network, we computed all the cliques using NetworkX (Hagberg *et al.* (2008)). Cliques are a concept in network theory that describe groups of nodes in a network that are fully connected among themselves. This is relevant because “in the context of coevolution, a clique represents a set of residues wherein each residue covaries with all of the others” Lee *et al.* (2012). We then produced a square scoring matrix by matching the cliques of the residue-residue coevolution networks e.g. if a clique in enzyme A has X coevolving pairs, and all X coevolving pairs have ECPs in enzyme B, then they are Equivalent Coevolving Cliques (ECC), and we increment the score between A and B.

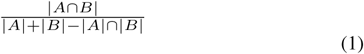

The clique similarity scores were then normalised to values between 0 and 1 by turning them into Jaccard distance scores where A and B are the cliques of the residue-residue coevolution networks for enzymes A and B (Equation 1): a score of 0 means no similarity between the networks, while a score of 1 is an exact similarity between the networks. From these scores, we produce CSNs where nodes are enzymes and edges are made between two nodes if their similarity value is above some arbitrary threshold.

### 2.7 Enzyme Substrate Specificity Prediction Through Label Propagation

To test the predictive power of the networks in labeling unknown data, and to analyse how well the networks produced capture the diversity in substrate specificity of the enzymes in the datasets, we perform a label propagation experiment on the EC labels of the datasets. We implemented an algorithm inspired by the work of (Schwikowski *et al.* (2000)) in their functional prediction of the yeast protein-protein interaction network. The following pseudocode was used:

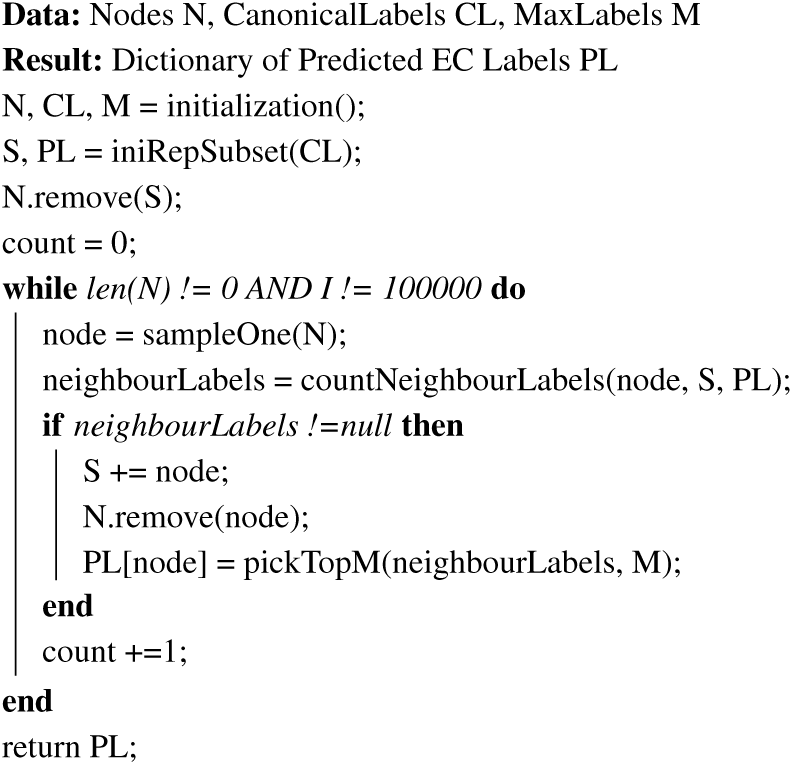

The algorithm starts by initialising four variables: the list of nodes N of some network (either an SSN or a CSN in this case), a dictionary CL of the canonical EC number annotation taken from Swissprot (one or more EC number per enzyme of the network), and a max number M of predicted labels per enzyme. The algorithm outputs a dictionary PL that will contain the annotation predicted by the algorithm, where keys are entries and values are the labels.

Unlike the approach described by Schwikowski and co-workers (Schwikowski *et al.* (2000)), an initial semi-random subset S of nodes is selected, and the dictionary PL with the labels for those nodes is initialised. S is a representative subset that contains at least one enzyme for each EC number. The algorithm then remove the nodes of S from N, and then progressively predicts annotations for the rest of the nodes through label propagation.

After initialisation, the algorithm iterates through the remaining nodes one at a time in random order, ranking the labels of a node’s neighbours based on propensity of the EC number. If the node has annotated neighbours by that iteration, the node is added to S, is removed from N, and the top M EC numbers are assigned to the node in dictionary PL. This is repeated until the size of N reaches zero, or until a maximum amount of iterations is reached. As this algorithm is stochastic, it is necessary to perform multiple iterations.

In order to determine the predictive power of this approach when applied to both SSNs and CSNs, we needed to iterate through the possible thresholds for both networks. This would let us identify the optimal threshold for each dataset for each network type, therefore allowing an in-depth comparison of the two methods based on currently known optimal conditions.

Specifically, for each threshold ranging from 0.05 to 0.9, we performed 200 iterations of the label propagation algorithm for a MaxLabel value of 2 for both network types. For each threshold, we produced two metrics: average precision and average recall over the 200 iterations. In our case, true positives are when the correct annotation is predicted, false positives are when an incorrect annotation is predicted, and false negatives are when no annotation is predicted. Specifically, the precision and recall values produced only take into account nodes not included in the initial representative subset, and nodes which have complete EC numbers.

It is important to consider what these two metrics represent in terms of network structure and available labels. Depending on the network structure drawn by the threshold, a different set of nodes can get labels propagated to them, and the precision only assesses how correctly the labels were predicted for that set, no matter how large or small it is. The recall on the other hand will describe the extent the network could propagate labels, no matter how correct those propagated labels are. While the best network will optimise both values, the precision should be handled with caution due to its larger dependence on the correct labels being known. For example, a correctly connected but incorrectly labeled promiscuous component will penalise the precision after label propagation. The recall is less affected by this problem, as it is more dependent on the overall network structure rather than the labels in this case. It is able to assess how well a particular network covers the enzymes, as it would be heavily penalised for leaving nodes as edgeless.

Therefore, while a balance of both precision and recall is necessary, more emphasis is given to the recall when picking the optimal threshold. We plotted the results of the label propagation onto Precision-Recall curves (Figure 1 in Supplementary Data), and identified the optimal threshold for SSNs and CSNs from both families by ranking the thresholds by sorting by F1 score, which is the harmonic mean of both the precision and recall, and then selecting the threshold with the highest recall.

**Fig. 1.**
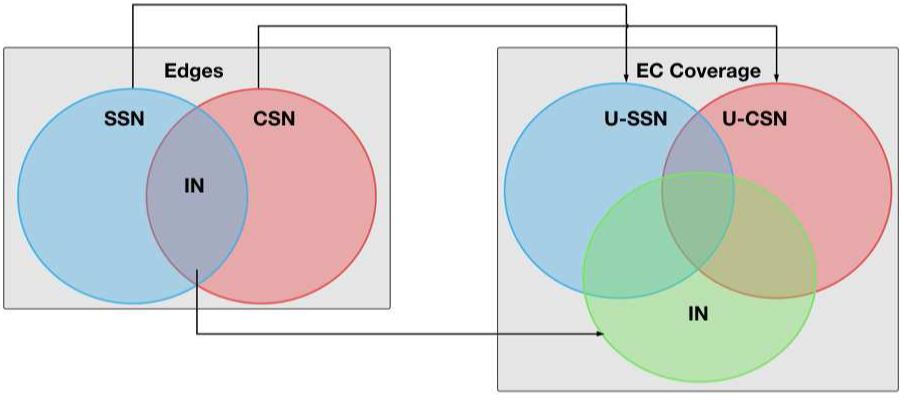
Venn diagrams explaining the comparison of EC coverage between the SSN and CSN. IN stands for Intersection Network, U-SSN for Unique Sequence Similarity Network, and U-CSN for Unique Coevolution Similarity Network

### 2.8 SSN and CSN Comparison

Metrics were computed to compare the topology of SSNs and CSNs, including the number of shared edges, number of network components (i.e. independent subgraphs) and the number of edgeless nodes. The network comparisons were carried out at the optimal predictive threshold for both SSNs and CSNs as determined above. The networks were visualised using Cytoscape and laid out using the organic layout algorithm (Shannon *et al.* (2003)). The networks can be seen in Figures 2 and 3.

**Fig. 2.**
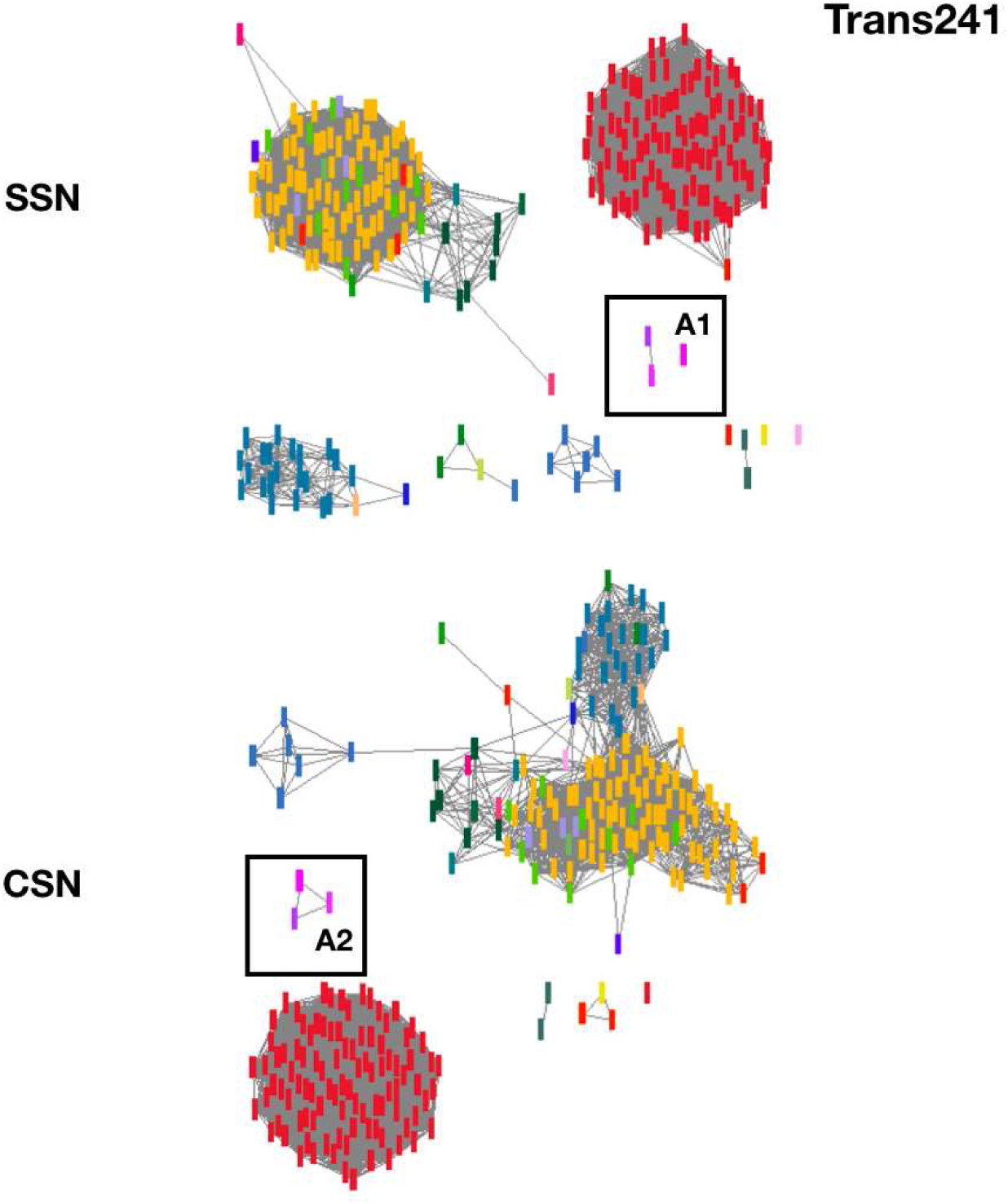
Visualisation of the optimal SSNs and CSNs for the Trans241 dataset. The top network is the SSN at 0.34 threshold and the bottom left network is the CSN at 0.30 threshold. Nodes are coloured based on the EC annotation from Swissprot. The A1 and A2 boxes represent the neamine transaminases for the SSN and CSN respectively.

**Fig. 3.**
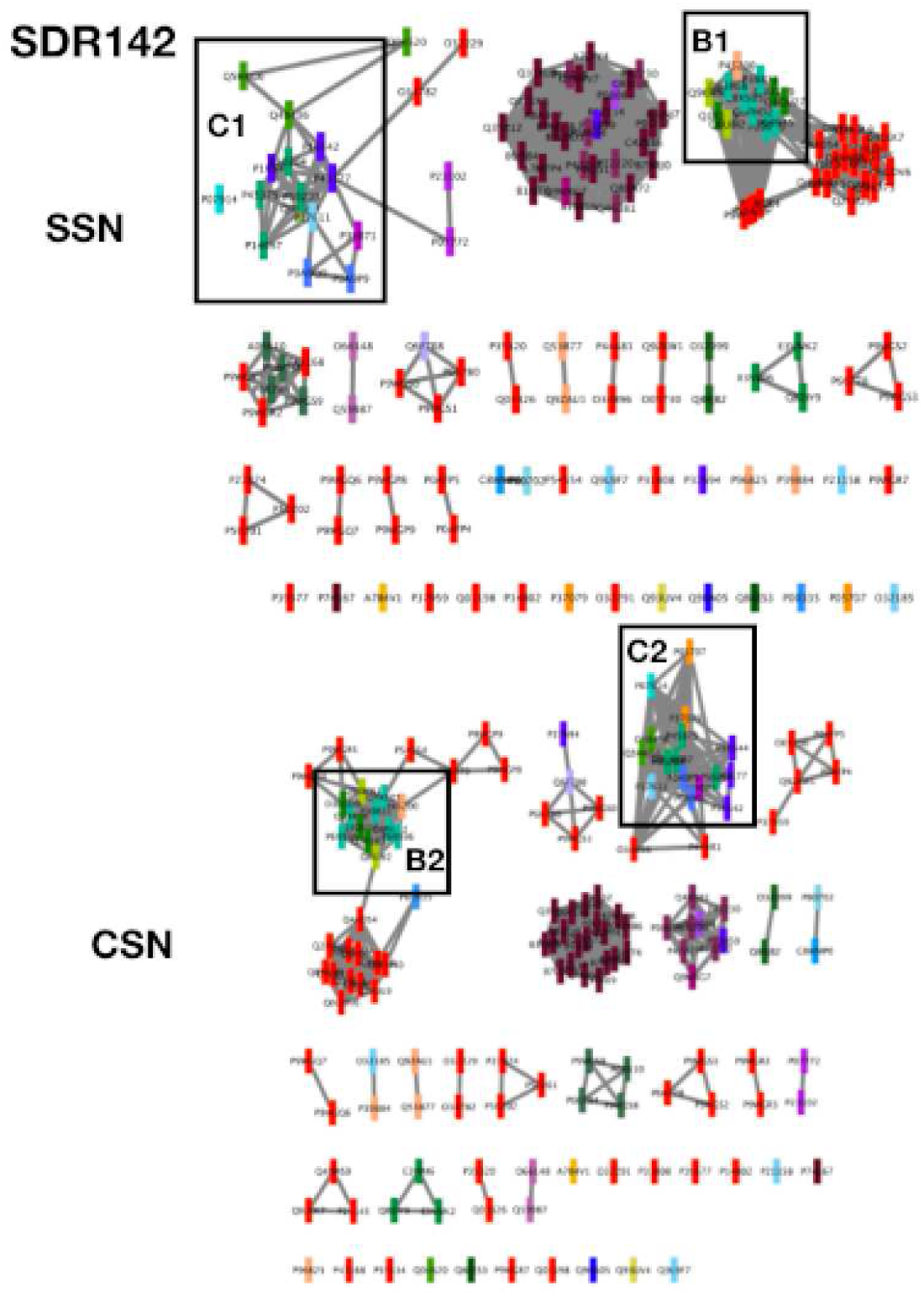
Visualisation of the optimal SSNs and CSNs for the SDR142 dataset. The top network is the SSN at 0.33 threshold, the bottom network is the CSN at 0.45 threshold. Nodes are coloured based on the EC annotation from Swissprot. The B1 and B2 boxes represent the first promiscuous component discussed for the SSN and CSN respectively. The C1 and C2 boxes represent the second promiscuous component discussed for the SSN and CSN respectively.

#### 2.8.1 SSN and CSN Structural Comparison

The set of edges that are shared by both a SSN and CSN we call Intersection Network (IN) was computed by overlaying the networks and identifying common edges between the networks. We also create networks for the edges unique to the SSN and CSN respectively, which we call Unique-SSN (U-SSN) and Unique-CSN (U-CSN) (Fig. 1). For each of these networks, we compute the number of EC labels covered. For the U-SSN and U-CSNs, we also compute the more specific number of EC labels completely unique to them, by disregarding EC labels also present in the IN and the counterpart unique network.

#### 2.8.2 SSN and CSN Functional Comparison

Different network components can have multiple protein clusters comprising enzymes of different substrate specificities. It is important to be able to quantify the distribution of EC labels across clusters in the network. To do this we computed a metric that we refer to as the MaxClust. The MaxClust is the largest single network component that contains nodes for a given EC number. More specifically, we computed the size of the MaxClust for every EC label, indicating substrate specificity, (called MCNumber), and then we computed the fraction of nodes of that EC label covered by the MaxClust (called MCFraction). From these two values, we calculate a metric we call Weighted-Average MaxClust Coverage, or WAMCC. The WAMCC is calculated for a network and represents how well connected enzymes of a similar substrate specificity are in a given network. While this metric does not give details about individual specificity classes across components, it works as a good comparison metric to see how specificities differ across the network and how much the two network types agree on the general network topology.

WAMCC is calculated as follows:

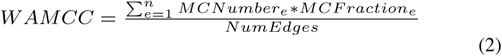

To perform a comparative analysis between CSNs and SSNs, we also produce SSNs for both the Trans241 and SDR142 datasets, using the optimal sequence similarity threshold as determined by the precision-recall analysis described above.

## 3 Results

SSNs have been shown to be valuable for the functional analysis and annotation of enzymes and visual analysis of protein families to reveal themes and outlines (Atkinson *et al.* (2009)). Here, we consider the properties of CSNs with respect to each of these applications. For functional annotation we retain a particular focus on the assignment of enzyme substrate specificity.

### 3.1 SSNs and CSNs for the prediction and analysis of enzyme substrate specificity

#### 3.1.1 Computing the optimal threshold for the SSNs and CSNs

As described in the methods section 2.6 optimal thresholds were computed on a balance of precision and recall. Figure 1 in the Supplementary Data shows the precision recall graphs for the SSNs and CSNs from each family with the thresholds indicated.

For the Trans241 dataset, the thresholds are 0.34 and 0.30 for the SSN and CSN, respectively. For the SDR142 dataset, the thresholds are 0.33 and 0.45 for the SSN and CSN, respectively. The networks with these thresholds were then considered to be the optimally predictive SSN and CSN for both families.

#### 3.1.2 Comparison of the substrate annotation power of the SSNs and CSNs

In order to compare the utility of the SSNs and CSNs for the prediction and subsequent annotation of enzyme function, especially substrate specificity, we applied a label propagation approach to the optimally predictive networks identified above to recover the known specificities of the two families well characterised enzymes under study.

As can be seen in Table 1, we can see that for the Trans241 dataset, the SSN has a slightly higher precision, and the CSN a slightly higher recall. Whilst these difference are small for the network as a whole, they appear to be significant when the network is considered in detail. We notice that in some cases the SSN fails to assign substrate specificities where the CSN succeeds. For the optimally predicted thresholds there are no cases where the SSN is able to predict substrates that were not identified by the CSN.

**Table 1.** Label Propagation experiment results for the ooptimal SSN and CSN for both the Trans241 and SDR142 datasets. To produce the precision and recall metrics, we consider that true positives are when the correct annotation is predicted, false positives are when an incorrect annotation is predicted, and false negatives are when no annotation is predicted.

| Dataset | Trans241 |  | SDR142 |  |
| --- | --- | --- | --- | --- |
| Network (Threshold) | SSN (0.34) | CSN (0.3) | SSN (0.33) | CSN (0.45) |
| Precision | 0.934 | 0.918 | 0.725 | 0.753 |
| Recall | 0.987 | 0.995 | 0.973 | 0.982 |

For example, we can see in Table 3, for the Trans241 dataset, that the U-CSN, a smaller network than the U-SSN, includes two EC labelss that are unique to it, while the U-SSN includes no unique annotations. These EC numbers are 4.1.1.64 and 2.6.1.111 and in the CSN are assigned to two proteins (P16932 and B0VH76, respectively) one for each EC number, but these protein nodes are edgeless in the SSN. We also notice that of the neamine transaminases (2.6.1.93), of which there are three in Trans241, only two are connected into a single component in the SSN, while in the CSN all three are fully connected into a single component(A1, A2, Figure 2). Whilst the the sequence similarity threshold could be lowered to a point to add these connections to the SSN, it is necessary to lower the threshold to 29%, which is highly detrimental to the overall predictive accuracy of the SSN through the introduction of noise in the form of false positive predictions. This phenomenon shown by the significant reduction in label propagation precision from 0.934 to 0.851 at this reduced threshold (Figure 1 in Supplementary Data). These results indicate, for this protein family, for the optimal CSN and SSN threshold, the CSN is able to cover more of the substrate specificity distribution of the Trans241 dataset without having to sacrifice better precision.

**Table 3.** Functional topology metrics for the optimal SSN and CSN for both the Trans241 and SDR142 datasets. The metrics were also produced for the network induced by the edges the compared SSN and CSN pairs share, which we call Intersection Network (IN), and for the edges unique to each network, which we call Unique-SSN and Unique-CSN (U-SSN and U-CSN).

|  | Trans241 |  |  | SDR142 |  |  |
| --- | --- | --- | --- | --- | --- | --- |
|  | Edges | Covered ECs (Unique) | WAMCC | Edges | Covered ECs (Unique) | WAMCC |
| SSN | 8,605 | 24 | 97.915 | 725 | 31 | 58.564 |
| CSN | 6,677 | 24 | 99.174 | 537 | 31 | 57.955 |
| IN | 6,343 | 18 | 98.19 | 429 | 23 | 60.783 |
| U-SSN | 2262 | 16 (0) | 96.16 | 296 | 15 (0) | 71.508 |
| U-CSN | 334 | 22 (2) | 68.147 | 108 | 15 (3) | 45.44 |

For the SDR142 dataset, the optimal CSN has a higher precision and recall than the SSN as shown in table 3. The U-CSN, which is also smaller than the U-SSN for this family, includes three EC labelss that are unique to the U-CSN, while the U-SSN includes no unique annotations. These annotations are 1.1.1.140, 1.1.1.395, and 1.1.1.56, applied to four proteins (P05707 and P37079, P07914, and P00335, respectively) in the U-CSN. These proteins are edgeless in the SSN and therefore not annotated. Lowering the SSN sequence similarity threshold to 30% results in these enzyme classes are no longer edgeless, but doing so lowers the precision from 0.725 to 0.62. While more coverage is important, such a reduction in precision is an indication of noise being introduced, as the label propagation algorithm is less able to correctly assign specificity. These data therefore show that the optimal CSN is also a better predictive structure than the optimal SSN for the SDR142 dataset.

### 3.2 A comparison of general network and functional topology between SSNs and CSNs

SSNs and CSNs were produced for both the Trans241 and the SDR142 datasets at thresholds optimal for predictive power as described in the methods. Table 2 shows some general network statistics for these networks. CSNs equivalent to SSNs, in terms of maximum number of shared edges and other networks statistics, as described in the Methods, were also produced.

**Table 2.**
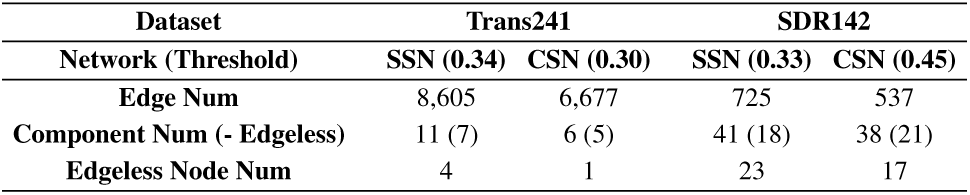
General network metrics for the optimal SSN and CSN for both the Trans241 and SDR142 datasets.

The first row of Table 2, shows the numbers of edges in each network. The optimal SSNs have significantly more edges than the CSN for both datasets. The second row contains the number of components (independent subgraphs) in each network. As edgeless nodes technically count as individual components, but are uninformative, we subtracted the number of edgeless nodes. The resulting values are shown in parentheses. Disregarding edgeless nodes, both network possess a similar number of components indicating that while the SSN and CSN may differ in certain details, they agree on a core of overall network structures and node distribution. Finally, the last row of table 2 shows the number of edgeless nodes in each network. In each comparison, across both datasets, the CSN has significantly fewer edgeless nodes than the SSN. As the optimal CSN has significantly less edges than the SSN, the fact that it also has fewer edgeless nodes could demonstrate a stronger robustness to change of thresholds.

Table 3 shows the distribution of EC coverage across the components in the network as measured by the WAMCC value. The WAMCC value was computed for both the SSN and CSN for both datasets. It is also computed for the different Intersection Networks and Unique Networks. Table 3 also contains the number of EC labels covered by all the aforementioned networks, including the number of EC labels unique to the Unique Networks. The SSN and CSN agree on the functional distribution of EC labels to a large extent. For example, the SSN and CSN of the Trans241 dataset have high WAMCC values of 97.915 and 99.174, respectively. Their intersection network contains 18 of the total 24 EC labels, which also represents agreement in overall functional topology. Similarly for the SDR142 networks, the WAMCC values are 58.564 and 57.955 for the SSN and CSN, respectvely. Also, 23 of the total 31 EC labels exist in the intersection network.

### 3.3 Complementary Analysis of Potential Promiscuity using SSNs and CSNs

Enzyme promiscuity is mostly unreported on the sequence databases. For example, most enzymes in SwissProt are annotated with only one EC number. However, we can still estimate the ability of both network types to identify potentially novel promiscuous clusters of enzymes. The simplest indication of potential promiscuity revealed by the networks is a connection between two enzymes with different EC annotation. In Figure 3, we can see that the SSN and CSN of the SDR142 dataset share a component (B1 and B2) that is very diverse in EC labels, with four different EC numbers shared across the 12 nodes, and one incomplete EC (B1 and B2). To further analyse this component, we examined three specific enzymes from this component that are annotated with all four EC numbers: Q1R183 (1.1.1.313), Q8U8I2 (1.1.1.276), and P39831 (1.1.1.298;1.1.1.381).

One potential reason for the connection of these three enzymes could be conservation of their function, increasing the chances of shared substrates, and therefore promiscuity. We used a tool called EC-BLAST (Rahman *et al.* (2014)) to compare the reactions catalysed by these three enzymes, namely by looking at bond changes and similarity in reaction centers. EC-BLAST returns the top 100 matches, along with similarity scores (Tanimoto coefficient) and a p-value. While the EC number 1.1.1.381 unfortunately could not be found on the database, the other three could. Indeed, the reactions 1.1.1.313, 1.1.1.276, and 1.1.1.298 were found in each other’s top 100 EC-BLAST matches, with the highest p-value being 1.4E-02. The similarity scores were 0.76 (1.1.1.313-1.1.1.276), 0.76 (1.1.1.313-1.1.1.298), and 0.89 (1.1.1.276-1.1.1.298). Given these scores, the amount of evidence showing that edges in both network types correlate with similar function, and the fact that both the SSN and CSN report these connections, it could be an indication that these three enzymes are promiscuous and share much of the reported catalytic diversity.

Another example of potentially novel enzyme promiscuity is the set of nodes indicated by C1 and C2 in Figure 3, the majority of which are connected in both the SSN and CSN. Much like the previous example, this component has a high EC number annotation diversity. Three enzymes in this component are of particular interest: P14697 (1.1.1.36), P07914 (1.1.1.395), and Q9X6U2 (1.1.1.30).P14697 and P07914 are both reported to catalyse CoA conjugated substrates: (3R)-hydroxybutanoyl-CoA (1.1.1.36), and CoA conjugated 3-hydroxy bile acids (1.1.1.395), respectively. 1.1.1.395 is unfortunately not available on EC-BLAST and so reaction similarity was not available for P07914. Querying 1.1.1.30 with EC-BLAST showed that 1.1.1.36 is the highest scored match, with a similarity score for bond changes and reaction centers of 0.98 and a highly significant p-value of 9.4E-04. Moreover, the substrate catalysed by Q9X6U2, (R)-3-hydroxybutanoate, is a similar substrate catalysed by P14697, (3R)-hydroxybutanoyl-CoA (1.1.1.36), difference by the CoA molecule. These analyses provide strong indications of potential promiscuity between Q9X6U2 and P14697,together with a potential relationship between P07914 and P14967 through the conjugation of CoA. As all three enzymes are part of the same diverse cluster, we can hypothesise that the relation between the three enzymes is transitive, resulting in Q9X6U2 potentially sharing catalytic activity with P07914. Importantly, while P07914 and its EC annotation of 1.1.1.395 is part of the component in the optimal CSN, as mentioned in section 3.2, this enzyme is edgeless in the SSN. While experimental data is required to validate these predictions, it is nonetheless an interesting result.

## 4 Discussion and Conclusion

SSNs are increasingly finding utility for the visual inspection and function analysis of protein families. The depiction of the functional relationships between proteins in a family as a network allows the visualisation and analysis of trends and groupings within large families (Atkinson *et al.* (2009); Gerlt *et al.* (2015)). Moreover, representation of protein families as networks makes their data accessible to common graph-based analytical approaches, metrics and tools, such as cluster analysis. SSNs have been widely applied to the functional analysis of enzyme protein families where they can help resolve substructure in the family and be used to assert functional equivalence.

Here we introduce a new approach for building networks of enzyme protein families, coevolution similarity networks (CSNs), that also depicts family structure and functional relationships but is not based directly on sequence similarity. We propose that since residues that coevolve do so under evolutionary pressure to maintain stability of structure and function (Hopf and Marks (2017)), a similarity measure computed on such residues should provide additional information about the conservation of protein function to that of global and local sequence similarity. SSNs have previously been shown to be valuable for functional annotation and protein family assignment using guilt by association type approaches. In order to investigate the utility of CSNs for the annotation of enzyme function, we evaluated the predictive power of CSNs for the re-annotation of unlabeled nodes in a partially annotated dataset. A label propagation algorithm was used to compare the predictive power of the SSN and CSN in assigning EC numbers to nodes whose labels were removed for the sake of testing, given a representative subset. In order to carry out a valid comparison between our SSNs and CSNs it was necessary to devise a reproducible approach to computing the weighting threshold for the removal of edges in the network that resulted in the optimally predictive network. We showed that the CSNs overall perform bettwe at SSNs for grouping together enzymes of similar substrate specificity. In summary, while the SSNs and CSNs are in agreement in a core of the overall network topology, we observed that the optimised CSNs are more predictive of enzyme substrates than optimised SSNs.

While CSNs contain novel enzyme functional information, they are very much complementary to their sequence similarity counterpart. First, both types of networks agree on large portions of the functional landscape that is predicted for a set of enzymes. Second, both network types comprise nodes which indicate proteins, integrating these two sources of information is as easy as combining their edges into one network. Finally, while the lack of promiscuity annotation is a problem that goes beyond the comparison of the SSN and CSN, identification of promiscuity is something the two network types can compliment each other on. While the latter does seem more apt for taking into account promiscuity when connecting enzymes, potentially promiscuous clusters appearing in both network types can give us more confidence that some cluster is potentially promiscuous. While we cannot be certain of some of the assumptions made without experimental data, new avenues for research and identification of promiscuity open through the CSN’s ability to make these interesting novel connections.

One of the heavy disadvantages of the CSN is that they are less accessible than SSNs - they are computationally more complex to produce, and coevolution is a harder metric to interpret directly. Also, CSNs are a more parametrised network type compared to SSNs. For example, the residue-residue coevolution networks are built based on picking the top N pairs most likely to coevolve, where N is currently just an arbitrary parameter. The filtering step where we disregard residue-residue coevolution pairs that are common to 60% of enzymes is also an arbitrary parameter. Optimising these parameters could therefore potentially improve the method as a whole. Also, the SSNs we produced in this work are based on pairwise global alignments, while other works have used metrics like BLAST similarity (Atkinson *et al.* (2009)) for producing SSNs.

While this work has focused on substrate specificity on the family level, future work includes applying CSNs to a metagenome to distinguish between families in general. Overall, while CSNs seemingly have an advantage over SSNs for relaying the functional distribution of a set of enzymes, they can could be used together to perform stronger analyses of enzyme sets.

## Supporting information

All the supplementary data mentioned in the manuscript

