## Supplementary material for "Functional Analysis of Enzyme Families Using Residue-Residue Coevolution Similarity Networks": All the supplementary data mentioned in the manuscript

**Trans241 Entries:**

P22256, P12995, P9WQ81, Q9I700, P40732, P50457, P53555, P18335, H8WR05, O30508, P77581, P23893, P42588, P24630, Q93R93, P9WPZ7, B0VH76, Q9APM5, Q7M181, Q53U08, O52250, P59324, Q88RB9, P22805, P38021, M1GRN3, Q5SHH5, P16932, Q8X4S6, P59317, Q8Z1Z3, Q9I6J2, O66442, P59321, P28269, P46395, P57600, Q7N9E5, P59086, P53656, P36568, P94427, Q01767, Q55665, Q6L741, Q9I6M4, Q9X2A5, E1V7V7, P12677, P0A4X7, O25627, O66557, Q74CT9, P45488, P44426, Q2FVJ6, Q9FCC2, P9WQ79, B7GHM5, Q9ZKM5, P57379, Q89AK4, Q8K9P0, P9WQ80, Q31QJ2, A0QYS9, P40829, P9WMN9, P63505, Q8DLK8, Q9RW75, Q8D0D7, Q8YS26, A2BVE5, D2D3B2, P9WQ78, B1X023, Q3AWP4, Q7NN66, Q818W2, Q8ZPV2, P56744, Q9PIR7, Q4H4F5, Q7NPI4, A9WIS7, Q8KAQ7, B0JPW6, B2J7M9, Q81M98, Q7VDA1, A2BPW6, B3EI07, B4S3Q6, Q2JS70, A3PBK8, B3QSA6, Q2JMP7, A5GMT2, Q110Z9, A1BJG8, B7KA18, B4SGW1, B8G822, B3EKJ7, A2SKQ7, A2C0U2, A9BEA5, A7NKV1, Q3M3B9, Q7V2J3, A5UU40, Q3AP59, Q3B1A1, B8HYK1, B0CC57, A8G3J9, Q31C50, B3QRD2, A4SGT2, Q7VMS5, Q7VAS9, Q8CSG1, B1XIT5, B0TFV0, B7K2I1, Q7V677, Q7U598, Q0I8G1, A5GUJ2, Q3ALU9, Q7V0G0, P9WPZ6, Q46GT9, A2C7I7, P36839, Q7MAE6, B9LKS0, Q89VE9, Q7W7H6, Q885K0, Q89LG2, Q8Y6U4, Q9L1A4, P59320, Q8EHC8, Q9KU97, Q9AAL3, Q9KNW2, P73133, Q8P5Q4, P24087, Q9ZEU7, Q8X4V5, Q8FL16, P30949, Q725I1, Q9KLC2, P30900, Q9RWW0, Q9AP34, A0QR33, B7GIK0, B7GH35, Q2G283, A8ALD5, Q5ZVA6, Q2S1S3, A7MUU9, Q4L7G9, A4FPX3, Q7MHY9, Q1IWZ8, A5WC94, P0C2D9, B9DZG0, B2SKS0, A3NCF3, Q9CC12, Q9JYY4, P59316, Q5H3I5, Q8Y6J9, Q3JPN1, O66998, A3MMQ8, P46716, Q882K8, Q9CHD3, A1V1L0, P59322, Q8UI71, P59315, Q9K8V5, P63566, Q99T15, Q2FXR4, Q7VTJ7, Q9A652, Q97GH9, Q59282, O08321, Q98BB7, Q59928, Q8XWN8, P59319, Q87L20, Q7WKW5, Q8CUM9, Q9KEB0, Q9JRW9, Q8CRW7, Q9K8G3, Q9CNT1, Q828A3, Q7VSH3, Q8R7C1, Q92BC0, Q92SA0, P54752, Q9PDF2, Q05174, Q92AX5, Q6QUY9, Q8PH31, Q82UP3, Q8FTN2, Q5HN71, Q7WDN7, P63567, P59323, Q72RH8, Q7MH19, Q7U5R5, Q5HER0, P63569, Q9JTX9, Q7V8L1, Q5HP24, P48247, Q7A4T5

**SDR142 Entries:**

P14697, P39831, P80702, P07914, Q9RA05, Q8KES3, P9WGS9, Q9WXG7, Q48436, Q8RJB2, Q82IY9, Q9KWN1, P0CI31, Q1R183, P47227, P74167, P05707, P07772, Q1QU27, P0CI32, P0A9P9, Q93UV4, A0R610, A7B4V1, D3U1D9, P96825, P16544, Q46381, Q8XA72, P9WGP9, B7NRJ0, Q04520, B7N6C8, Q83QJ8, P39071, B7M7P4,

A7ZPY4, Q3YZ12, C4ZXB6, B1IVT6, B7LDD3, B1XB16, B6I5B4, Q0T1X8, B1LNJ7, A8A347, B5Z114, Q31XU9, P23102, P9WGS8, P66784, Q7N4V7, Q8FHD2, P9WGS3, P69936, P69935, Q8X505, Q83RE8, P72220, P08694, C8WMP0, Q59987, P45375, Q6F7B8, Q9L9F7, P47230, P08088, P50204, P50206, P37079, P17611, P50203, P50202, P50201, P27874, Q01198, P39577, E3VWK2, Q5HKG6, E3VWI6, O66148, P00335, Q9ZAU1, P0A9Q0, P37694, P31808, P43168, O32099, Q8U8I2, P9WGQ7, P13859, P9WGS1, P16542, P21158, Q9X6U2, P37959, P41177, P0AFP4, P9WGR5, O05730, P54554, O34782, O32291, P9WGS0, O32229, P14802, P25145, P66778, Q53877, Q9ZKW1, P9WGS2, P0AFP5, P9WGP8, P25970, P45200, P9WGR3, Q4L8Y1, Q99RF5, Q2FVD5, Q4A054, Q49WS9, Q8CN40, Q92EK7, P55434, P9WGR2, P9WGQ6, Q5HD73, P9WGR4, Q5HLD8, Q2FE21, P66780, Q7A3L9, Q8NUV9, Q6GDV6, Q6G6J1, Q03326, P35320, O34896, P9WGR7, P44481, P39884, O32185

Table S1: Table of EC Number changes made for this work

| **Enzyme** | **Old EC** | **New EC** |
|---|---|---|
| Q93R93 | 2.6.1.- | 2.6.1.LYS |
| Q5SHH5 | 2.6.1.- | 2.6.1.LYS |
| Q9RW75 | 2.6.1.- | 2.6.1.LYS |
| Q53U08 | 2.6.1.93;2.6.1.95 | 2.6.1.93 |
| Q6L741 | 2.6.1.93;2.6.1.94 | 2.6.1.93 |
| P80702 | 1.1.1.50 | 1.1.1.STEROID |
| C8WMP0 | 1.1.1.52 | 1.1.1.STEROID |

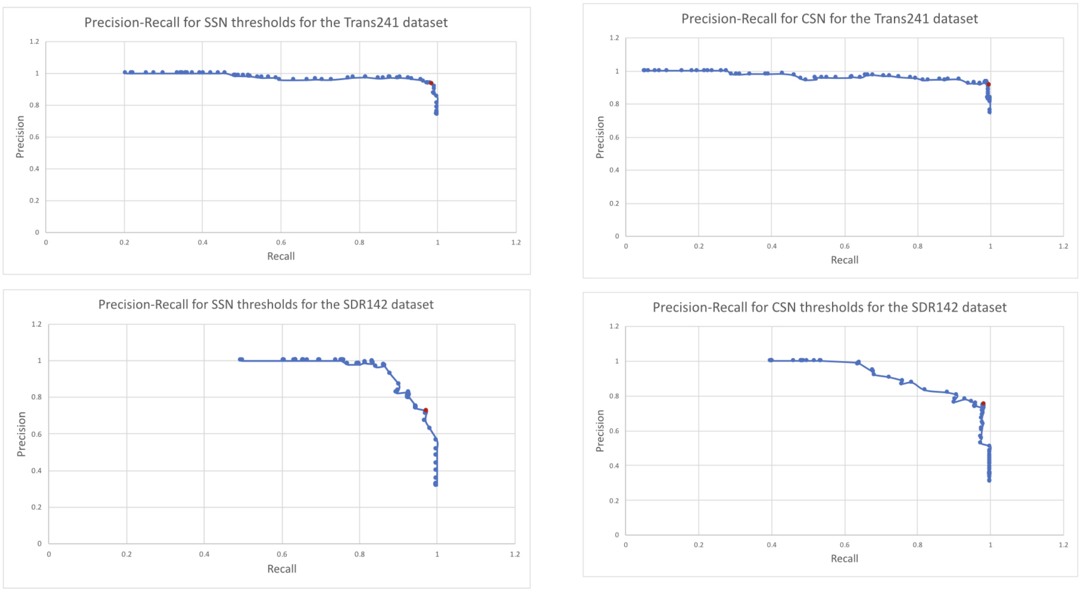


Figure 1: Precision-Recall curves for the optimised SSNs and CSNs for both the Trans241 and SDR142 datasets. The red point is the one selected.

Table S2: EC Number distribution for the Trans241 dataset

| **EC Number** | **Count** |
|---|---|
| 2.6.1.- | 2 |
| 2.6.1.LYS | 3 |
| 5.1.1.15 | 1 |
| 2.6.1.76 | 7 |
| 2.6.1.77 | 1 |
| 2.6.1.13 | 1 |
| 2.6.1.36 | 2 |
| 2.6.1.11 | 71 |
| 4.1.1.64 | 1 |
| 2.6.1.11; 2.6.1.81 | 1 |
| 2.6.1.18 | 2 |
| 2.6.1.19 | 1 |
| 2.6.1.105 | 1 |
| 2.6.1.93 | 3 |
| 2.6.1.11; 2.6.1.17 | 10 |
| 2.6.1.19; 2.6.1.22 | 6 |
| 2.6.1.48 | 2 |
| 5.1.1.21 | 1 |
| 5.4.3.8 | 98 |
| 2.6.1.62 | 21 |
| 2.6.1.81 | 3 |
| 2.6.1.82 | 1 |
| 2.6.1.111 | 1 |
| 2.6.1.113 | 1 |

Table S3: EC Number for the SDR142 dataset

| **EC Number** | **Count** |
|---|---|
| 1.1.1.333 | 4 |
| 1.2.1.n2 | 1 |
| 1.1.1.69 | 2 |
| 1.3.1.33 | 2 |
| 1.1.1.381; 1.1.1.298 | 6 |
| 1.3.1.19 | 1 |
| 1.3.1.56 | 6 |
| 1.1.1.395 | 1 |
| 1.3.1.- | 5 |
| 1.1.1.256 | 1 |
| 1.-.-.- | 53 |
| 1.1.1.276 | 2 |
| 1.1.1.313 | 3 |
| 1.1.1.56 | 1 |
| 5.1.3.34 | 1 |
| 1.1.1.30 | 1 |
| 1.1.1.- | 5 |
| 1.1.1.36 | 4 |
| 1.3.1.87 | 20 |
| -.-.-.- | 4 |
| 1.1.1.STEROID | 2 |
| 1.1.1.n4 | 1 |
| 1.3.1.25 | 2 |
| 1.3.1.28 | 1 |
| 1.3.1.49 | 1 |
| 1.1.1.201 | 1 |
| 1.1.1.340 | 3 |
| 1.1.1.325 | 1 |
| 1.1.1.320 | 2 |
| 1.1.1.304 | 3 |
| 1.1.1.140 | 2 |
